# Divergent macrophage induced lung pathology occurs between infants and adults during respiratory viral infection

**DOI:** 10.1101/2024.08.25.609597

**Authors:** David Verhoeven, Davis Verhoeven

## Abstract

Young children, especially those aged 4 months to 2 years of age, frequently exhibit severe morbidity during respiratory viral infections. For influenza infections, macrophages/monocytes serve as front line defenses against early viral replication in the lungs until the adaptive immune system arrives to clear virus. However, infiltrating inflammatory monocytes are a significant cause of influenza induced lung pathology/morbidity. We utilized a young murine model of respiratory viral infections using 21-day-old mice to investigate the mechanisms driving the heightened influenza induced morbidity observed in human young children. We hypothesized that macrophages/monocytes responses to influenza would diverge between young and older mice despite our evidence that macrophages from both groups appear to control viral replication at similar rates. While inflammatory monocyte infiltration contributed to influenza induced morbidity/lung inflammation in adults, they did not appear to contribute to morbidity in young mice. Instead, young mice appeared to develop lung inflammation through a lack of interferon gamma (IFNγ) and infection of macrophage populations. In contrast, adult mice controlled early viral replication through macrophage populations (alveolar, interstitial, and inflammatory) and inflammatory monocytes. While intrinsic limitations in anti-viral cytokine responses, especially IFNγ, characterized the macrophage response to viral infection in young mice, the innate immune response to infection appears diminished compared to adults. This study highlights the intrinsic limitations in macrophage effector functions that may arise in young children but that also contribute to disease pathology.

## Introduction

Respiratory viral infections especially affect the young and old and frequently cause primary or secondary pneumonia. Infants and toddlers (0.2-2 years old, referred here after as young children) are especially susceptible to influenza induced morbidity and frequently require hospitalization due to its severity or development of secondary bacterial pneumonia (1, 2). Young children undergo rapid immunological maturation and thus the immune responses of a young child can be quite different than one of neonates or older children. While host responses to influenza viruses have been well documented for neonates, which exhibit significant immunological limitations from low antigen presenting cell presentation (3, 4) and T-cell bias (3, 4), or older children, which have more comparable but not exact similarities to adults, less is known about how immunological maturation affects the young child’s response to influenza infection. Moreover, our knowledge about innate contributions to antiviral responses to infection in young children is limited especially with respect to resident/non-resident macrophage function in the lungs. Whether the enhanced morbidity in young children is a function of poor immunological control over viral replication, enhanced lung inflammation, or contributions from immunopathology are not well documented. Previous studies have clearly demonstrated that T-cell responses in young children, with severe respiratory viral infections, were limited (5, 6). Moreover, young children might exhibit prolonged viral replication in the lungs possibly due to limited adaptive immune responses (7). However, innate antiviral responses precede adaptive responses and thus the initial character of the early innate response could greatly influence the resulting level of lung pathology later in infection. Additionally, influenza virus gradually degrades the innate immune response so that the host becomes highly susceptible to secondary bacterial pneumonia around 6-7 days post infection (8–10). Given that children are likely to develop secondary bacterial infections (*e.g.* middle ear infections), influenza may have a bigger influence on macrophages of the young than older children/adults possibly indirectly through rampant IFNα production (10). We have previous determined that infant/toddlers may outgrow some of the limitations immunological maturation places on their CD4 T-cells. However, less is known about how (or even if) infants/toddlers outgrow limitations in their antiviral monocyte/macrophage functions. We also know little about macrophage contributions to the enhanced level of morbidity in young children given that children generally secrete less antiviral cytokines. Due to this, pathology through “cytokine storms” may be less likely as adults possibly due to other mechanisms.

Monocytes and macrophages in the lungs are a significant contributor to early control of viral replication (11) but also help shape the adaptive response. Specifically, alveolar macrophages (AvM) may contribute some antiviral responses but may play a role in protection from lung damage and respiratory failure (12, 13). In contrast, inflammatory monocytes and macrophages differentiating from monocytes (MDMo) or interstitial macrophages (IsMo) may play bigger roles in viral suppression and lung inflammation (14–16) with high pathologic strains capable of overcoming the antiviral responses of MDMo (17). IsMo cells are highly proliferative and secrete an abundance of IL-1 and IL-6 (18). AvMo can significantly down-regulate antigen presentation to T-cells (19, 20) as influenza infection in the young is characterized by poor T-cell responses (5, 6). In contrast, monocytes or MDMo may actively present antigen (directly or indirectly) to naïve T-cells in the draining lymph nodes (21). Various strains of influenza virus also have different propensities for macrophage infection and nonproductive viral replication. Moreover, human AvMo display 2,6α linked sialic acid moieties (22) which serve as the receptor for viral entry. Of interest, MDMo and resident alveolar macrophages have different propensities for viral replication with both cells readily infected but with only MDMo productively producing infectious virions (23). Furthermore, depending on the activation stimulus, macrophages can be programed into cells with pro-inflammatory characteristics (classical activation induced by IFN-γ) or anti- inflammatory/repair characteristics (alternative activation through IL-4) (24, 25). Thus, depending on the activation level, location, and possibly local viral production, the various macrophage populations in the lungs during influenza infection could have dramatic effects on lung pathology in the young as compared to adults.

To examine the effects of influenza on macrophages/monocytes with respect to the enhanced morbidity common in young children, we have developed a young murine model of infection. This age of mice exhibit many similarities to infant and toddler human children as stated in our other previous publications (26, 27). Here, we examined the contributions of infiltrating monocytes and macrophages to protection and immunopathology and found divergent responses between young and adult mice. We found that the character of monocyte and macrophage responses to infected epithelial depots was very different between both groups. Specifically, antiviral cytokines (*e.g.* IFNγ) were significantly lower in the lungs of young mice but tended to cluster near infected epithelial cells in the bronchioles young children and older children/adults. Despite effectively limiting viral replication similar to adult mice, the innate immune system of young children were clearly not as immunologically mature thus possibly contributing to the higher morbidity observed.

## Materials and Methods

### Mice

10 week old female BALB/c mice were purchased from the National Cancer Institute Biological Testing Branch and Jackson Mice (Jax, Bar Arbor MI) or BALB/c were bred and maintained under specific pathogen-free conditions and used at 21 days of age. CCR2 KO mice were obtained from Jax and bred and housed until 8-10 weeks or 21 days old with wildtype C57BL/6 used in similar ages. All animal Iowa State University (ISU) Institutional Animal Care and Use Committees. All mice, infected or not infected, were housed in biocontainment barrier rooms under ABSLII conditions. PulseOx readings were performed as done previously (28).

### Influenza virus infection

Influenza virus H1N1 PR8 (A/PR/8/34) were grown in the allantoic fluid of 10-day-old embryonated chicken eggs (Charles River, Wilmington MA) as previously described (29). Determination of influenza viral titers in 100mg lung homogenates were accomplished by the tissue culture infectious dose 50 (TCID_50_) assay as described previously (30). For *in vivo* infection, mice were anesthetized with isofluorane and 40 μl of influenza PR8 influenza virus containing 250 TCID_50_ or 500 TCID_50_ was administered intranasally respectively to young mice or adults. The difference in viral titers administered represents the 2-fold differences in lung weights and body weights between young and adult mice used in this study.

### Leukocyte cellular isolations

Lungs were digested with CTID media (collagenase II and IV, trypsin inhibitor, DNAse) overnight at 4 degrees. Cells were then pushed through a 100μM cell strainer and red blood cells lysed by incubation in ACK buffer. Cells were at least >85% viable after isolation as evidenced by trypan blue staining.

### Fibroblast isolations and infections

10cm^2^ lung tissue sections were harvested from uninfected infant and adult mice and cultured in DMEM 10%FBS using 24 well plates for seven days allowing fibroblasts to migrate from the lungs. Cells were then infected at MOI of 1 using influenza A/PR/8/34 reasssorted with PR8 for 1 hr (BEI resources, Manassas VA). Cells were then either fixed with 4% paraformaldehyde overnight after 30 hours of infection or fixed with RNAlater (Sigma) and stored overnight at 4 degrees after 48 hours of infection. For qRT-PCR, cells were harvested using RNeasy plus kit (Qiagen, Gaithersburg MD) and viral transcripts determined using primers for G3DPH (calibrator) or influenza M2 protein (Primer Bank). For fluorescent microscopy, cells were permeabilized with saponin buffer (Biolegend, San Diego CA) for 1hr and then stained with anti-influenza HA (1:1000 dilution, Thermofisher, Waltham MA) for 1hr. at room temperature followed by 5 washes of permeabilization buffer. Cells were then further incubated with anti-goat FITC (1:1000 dilution) in saponin buffer for 1hr followed again with 5 washes. Finally, cells were counter-stained with DAPI-gold (Vector labs, Burlingame CA) and imaged using an Olympus fluorescent microscope allowing software to optimize exposure for DAPI and FITC. Some fibroblasts were also placed into RNAlater (Sigma) for subsequent qRT-PCR.

### Abs and Reagent

We obtained CD45 FITC, CD11b PE, CD11c PercpCy5.5, DC pan-marker APC, IgM PeCy7, F4/80 ApcCy5.5, MHCII ApcCy7, DC pan-marker FITC (dump), CD3 FITC (dump), CD49d FITC (dump), and CD19 FITC (dump) from Biolegend (San Diego, CA). Cells were surface stained with staining with the fluorochrome-conjugated Abs as previously described (42). For purification of macrophages/monocytes, we obtained clone ER-HR3 from Biolegend (San Jose, CA) and used an anti-rat nanoparticle for purification (Polysciences, Warrington PA).

### Flow cytometry

Cells were acquired using an LSRII flow cytometer (BD, San Jose CA) with a minimum acquisition of 100,000 events. BrdU was determined after surface staining followed by antibody detection according to manufacturer’s directions (BD). Analysis of acquisition events was accomplished using FlowJo software (Tree Star, Ashland Oregon) with gating using live amine dye (Invitrogen, Carlsbad CA) and double discrimination.

### Cytokine assays

23-plex murine cytokine assays were performed by Luminex (Biorad, Hercules, CA) on lung homogenates from 50mg sections in cold PBS with 1% Igepals (Sigma) generated by repeated grinding with a pestle. Lungs were clarified by centrifugation at 10,000 RPM for 30 seconds.

Supernatants were transferred to new cold eppendorfs and snap frozen in liquid nitrogen.

### Histology

Mice were euthanized and perfused with 5ml of PBS by cardiac puncture after severing the dorsal aorta. Lungs were inflated with 4% buffered formalin by bronchiole infusion inflating the lung to 2x its size. Lungs were harvested and stored in 4% formalin. Fixed tissues were processed and stained with H&E (AML St Augustine FL or ISU pathology core facility 24hrs post-harvest). Tissues were blindly evaluated by a board licensed veterinary pathologist (ISU pathology core) similar to (31) and assigned a histopathologic score.

### RNAscope

10μm tissue lung sections were cut from 4 DPI infected mice or uninfected controls. Macrophages (F4/80) and IFNγ were stained using RNAscope probes commercially available (Advanced Cell Diagnostics, San Diego CA) and amplified according to the manufacturer’s directions. Tissue was then counter-stained using Hematoxylin/Eosin.

### qRT-PCR

RNA was extracted from tissues using RNeasy Plus (Qiagen). For the detection of changes in gene expression in influenza infected young and adult mice, the RNA levels for each were compared with the levels in uninfected young or adult mice (calibrators), and data are presented as the change in expression of each gene. The Δ*Ct* value for the tissue sample from the calibrator was then subtracted from the Δ*Ct* value of the corresponding lung tissue of infected mice (ΔΔ*Ct*). The increase in cytokine mRNA levels in lung tissue samples of the infected animals compared to tissue samples of baseline (calibrator) animals was then calculated as follows: increase = 2^ΔΔ*CT*^. Lungs and spleens were isolated from uninfected and 4, 7, and 10 days post-infected adult and young mice and stored in RNAlater (Sigma).

Superarray (Qiagen) qRT-PCR was performed per the manufacturer’s instructions using 260ng of RNA extracted from lungs (inflammatory cytokine and receptors panel) or spleens (T/B-cell activation panel). Data was analyzed using RT^2^ Profiler PCR Array analysis program to calculate calibrated fold changes (Qiagen). Primers for each were designed from PrimerBank for mouse and qRT-PCR was performed as a one step Sybr Green reaction using Luna kits (NEB, MA)

## Results

### Heightened morbidity during influenza infection of young mice

We previously determined optimal dosage of influenza for reliable infection of both young and adult mice based on lung weights similar to our prior studies (26, 27) and outlined in Fig 1A. Using weight loss as a marker of morbidity in these mice, we found young mice had significantly higher weight loss than adults (Fig 1B) during a sublethal infection (Fig 1B). We found that young and adult mice had similar viral titers at peak viral burden at 3 days post-infection but began to diverge by day 5 when the innate immune system is usually highly present in the lungs (Fig 1C). However, adults were clearing virus by 10 days post-infection contrasting with young mice that cleared virus more slowly.

**Figure 1.**
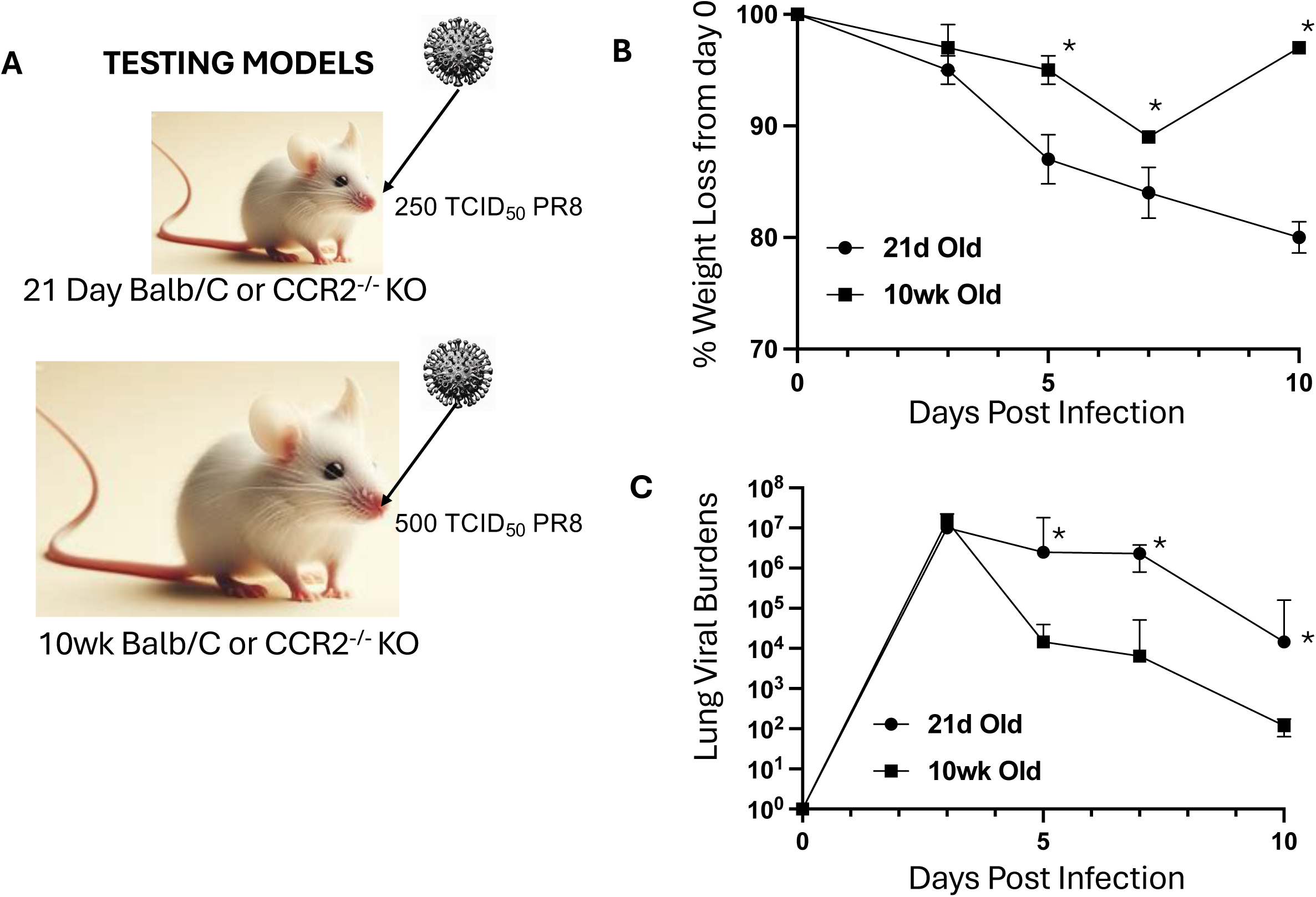
Young mice exhibit heightened morbidity during influenza infection despite similar early viral loads. (A) We infected mice with divergent lung doses as 21d old mice have approximately ½ the lung weight and size of 10wk old mice. We assessed **(B)** weight loss morbidity and **(C)** viral loads by TCID_50_ assay from whole lungs at the indicated time-points during infection. N=5 each for 2 reps. * is p- value <0.05.

We next photographed mice at day 5 post-infection and 21d old mice were having much higher levels of morbidity as evidenced my worse body conditions (Fig 2A). We also examined tissue histology at day 5 post infection and found that young mice exhibited divergent cellular infiltration at 5 days post- infection (Fig 2B) with areas of much higher lung consolidation.

**Figure 2.**
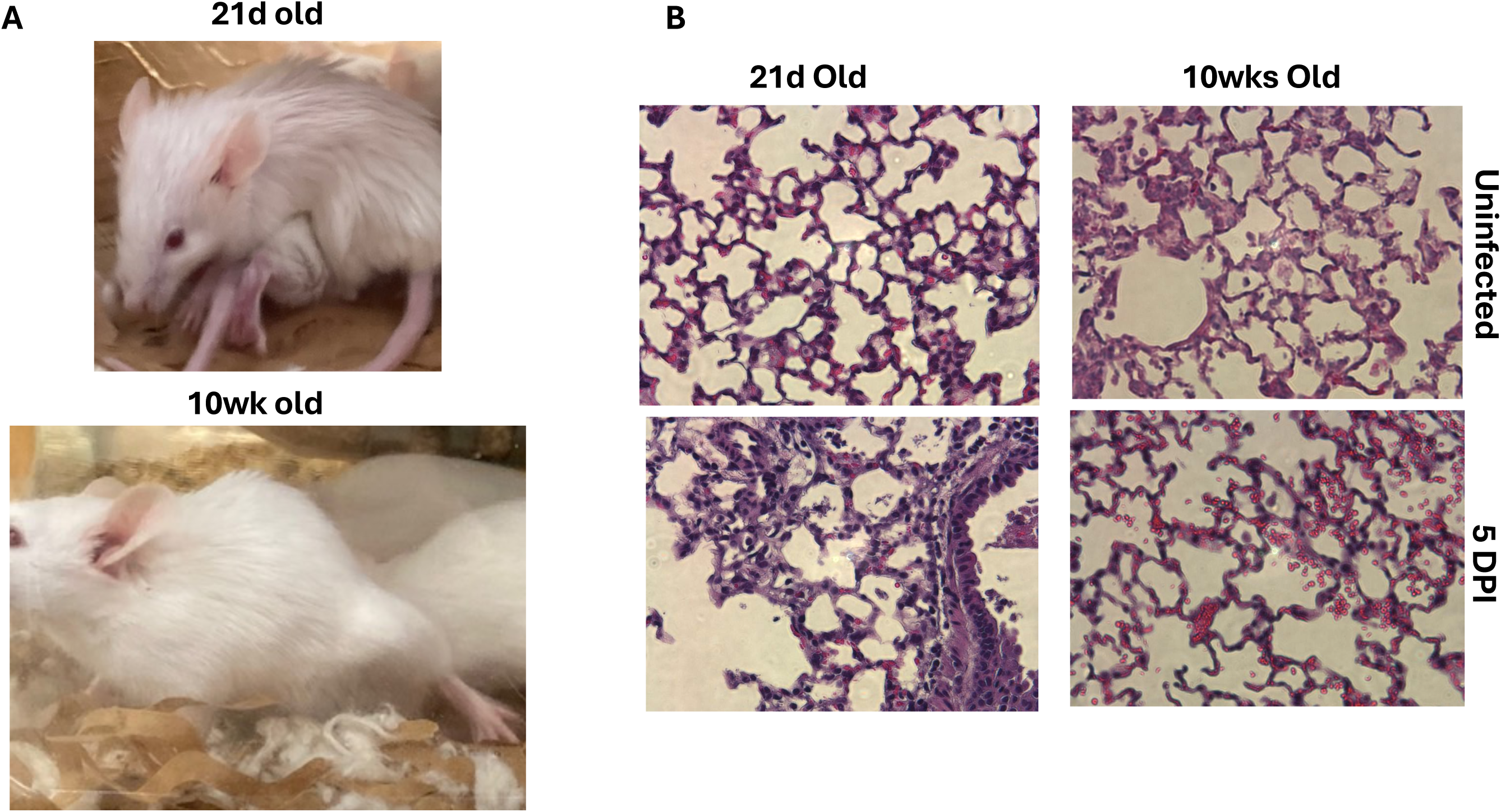
Body condition and lung histology both agree that younger mice have higher morbidity during infection. (**A)** We photographed the bodies of 21d and 10wk old mice to document their conditions at 5 days infection when virus in the lungs began to diverge. We did not capture it but differences in body condition were already beginning to appear as early as 3 days post-infection despite similar lung viral loads found in Fig 1. **(B)** Lung histology was obtained at day 5 post-infection as compared to controls.

### Inhibition of inflammatory monocyte infiltration only alleviates lung immunopathology in adults

Since viral burdens in the lungs were worse in 21d mice after infection and morbidity (body condition and weight loss) were worse at day 5, we wondered if high inflammatory monocytes/macrophages might be the cause of the higher morbidity. To examine the role of inflammatory monocytes/MDMo with respect to tissue pathology, we infected CCR2^-/-^ mice or C57BL/6 controls similar to our BALB/c mice using a sublethal dosage. Of interest, adult CCR2^-/-^ mice had less weight loss and significantly better lung histology than young mice (not shown). In significant contrast to 10wk old mice, 21 day old mice were not alleviated from lung histopathology if they could not recruit inflammatory monocytes after infection (Fig 3). This implied to us that other factors might be playing a role in failing to control viral replication to the same levels. We did access oxygen saturation in these mice and like their body conditions in shown in figure 2, 21d old mice has an average of pulseOx rate of 80% while 10wks were at 90% at day 5.

**Figure 3.**
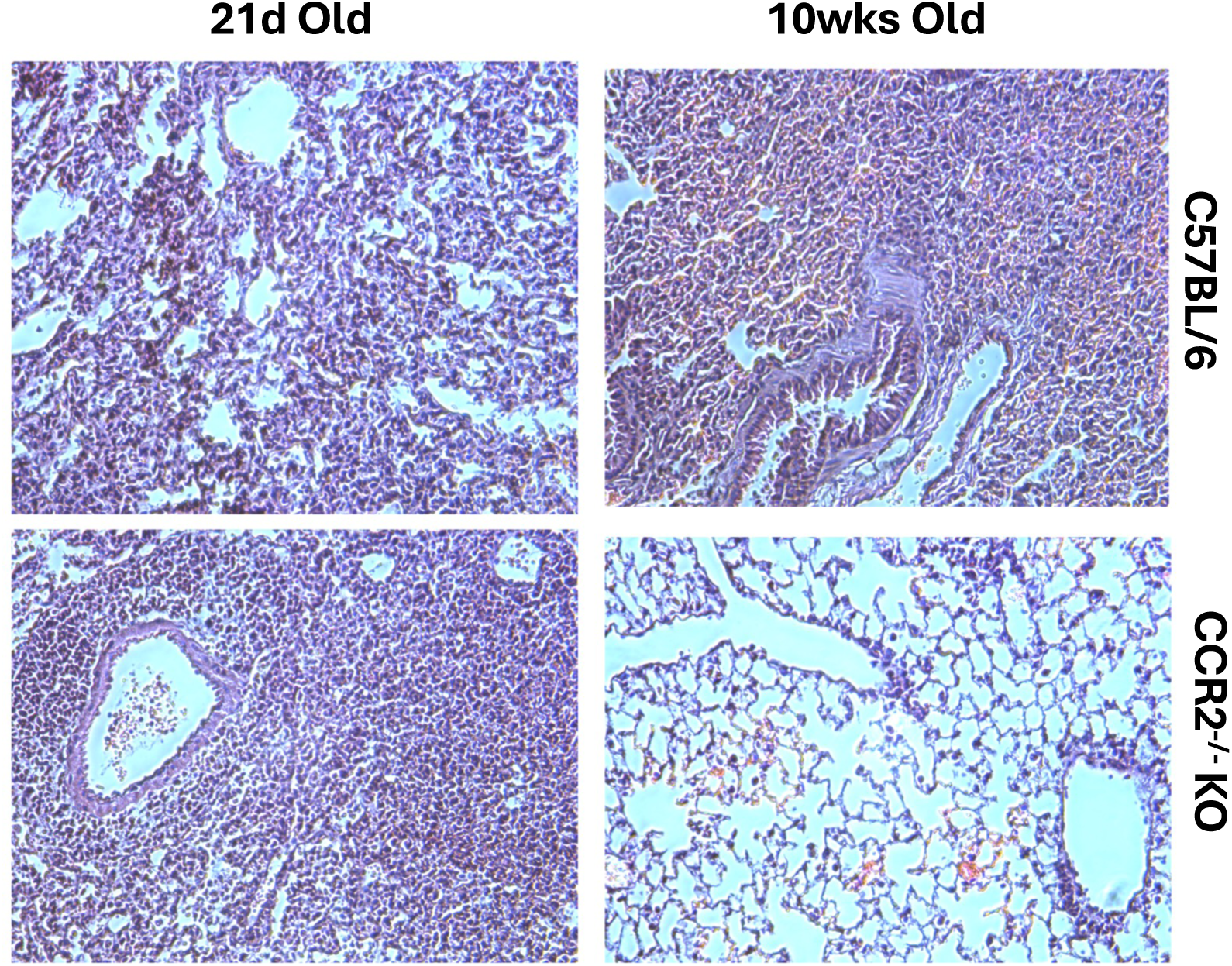
Limiting monocyte recruitment to the lungs only alleviates lung immunopathology in older mice. We infected C57BL/6 or CCR2^-/-^KO mice of the same ages as our BALB/c mice similar to fig 1A. We assessed the lung histopathology at day 5 post-infection in both groups of mice.

### Lungs have lower levels of IFNg in 21d old mice which corresponds to poor release from lung macrophages

Although not the sole source of IFNγ in lungs, macrophages are significant contributors of this cytokine. We assessed the cytokine levels in lung sections by Luminex and found the vast majority of cytokines did not diverge much expect for IFNγ, TNFα, and IL-17a which was much lower in 21d mice than 10wk olds (Fig 4A). We next examined the number of macrophages in the lungs at day 5 along with whether they stained for high levels of IFNγ by RNAscope. We found a high number of both in the lungs of 10wk old mice that was much lower in 21d old mice (Fig 4B). We next isolated total macrophages from the lungs of both groups of mice and compared them over their age matched uninfected controls. In agreement to Fig 4A-B, we found isolated macrophages had much lower IFNγ mRNA transcripts suggesting an impaired ability to release the cytokine (Fig 4C).

**Figure 4.**
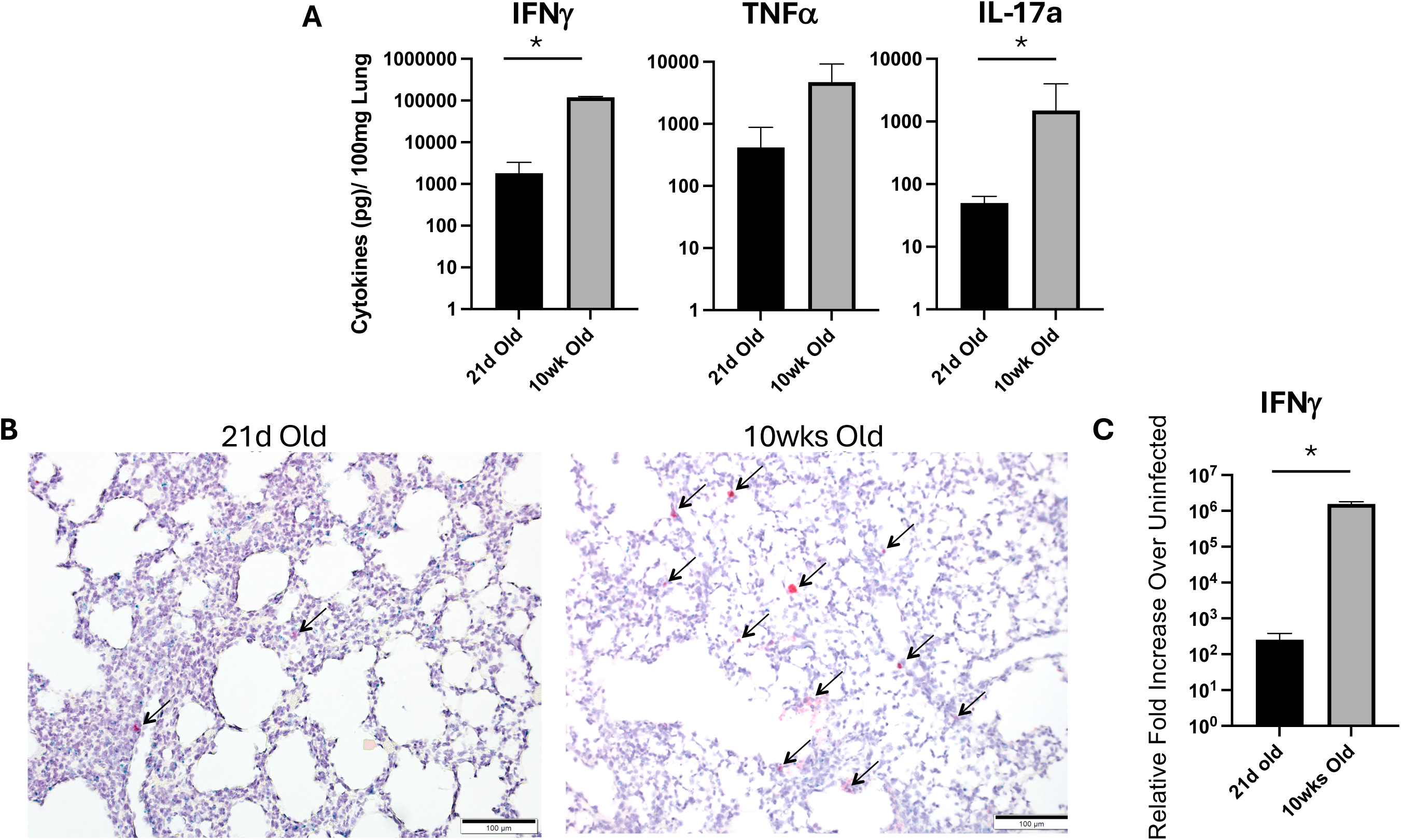
Lower IFNγ characterizes the lungs of young mice than older which appears to correlate with lower IFNγ+ macrophage. Lungs were obtained from uninfected and infected (5 days post infection) 21d old and 10wk old mice. **(A)** Levels of cytokines in ground/cleared lung homogenates were assessed by Luminex and only IFNγ, TNFα, and IL17a were highly expressed and different between the two infected age groups. Uninfected controls served as a baseline to subtract the levels of each cytokine found in the lungs. **(B)** RNAscope was used to probe for macrophages (F4/80) and IFNγ transcripts. Arrows point to macrophages that were positive for IFNγ transcripts at 5 days post-infection. **(C)** Macrophages/monocytes were isolated from the lungs of infected mice at 5 days post-infection and the level of IFNγ was confirmed in these isolated populations and expressed as increases over age matched uninfected controls. * p-value <0.05.

### Divergent control over IFNγ intracellular mediators occur in the macrophages from 21d old mice

We next evaluated whether there were any differences in macrophage activation (classical versus alternative) by qRT-PCR especially as IFNγ were so much lower in young mice. Of interest, macrophages isolated from 10wk old mice had significantly higher levels of STAT6 over controls but STAT1 dominated the response (Fig 5A-B). We found an increase in STAT1 expression, albeit trending lower than 10wk old mice, in younger mice. However, in contrast to adults, we found significant down- regulation of STAT3 and STAT6 over controls during infection in 21d old mice (Fig 5B-C). These data further suggest that diminished IFNγ in the lung of mice may alter the phenotype of antiviral macrophage populations and contribute to the dominance of classical activation in young mice.

**Figure 5.**
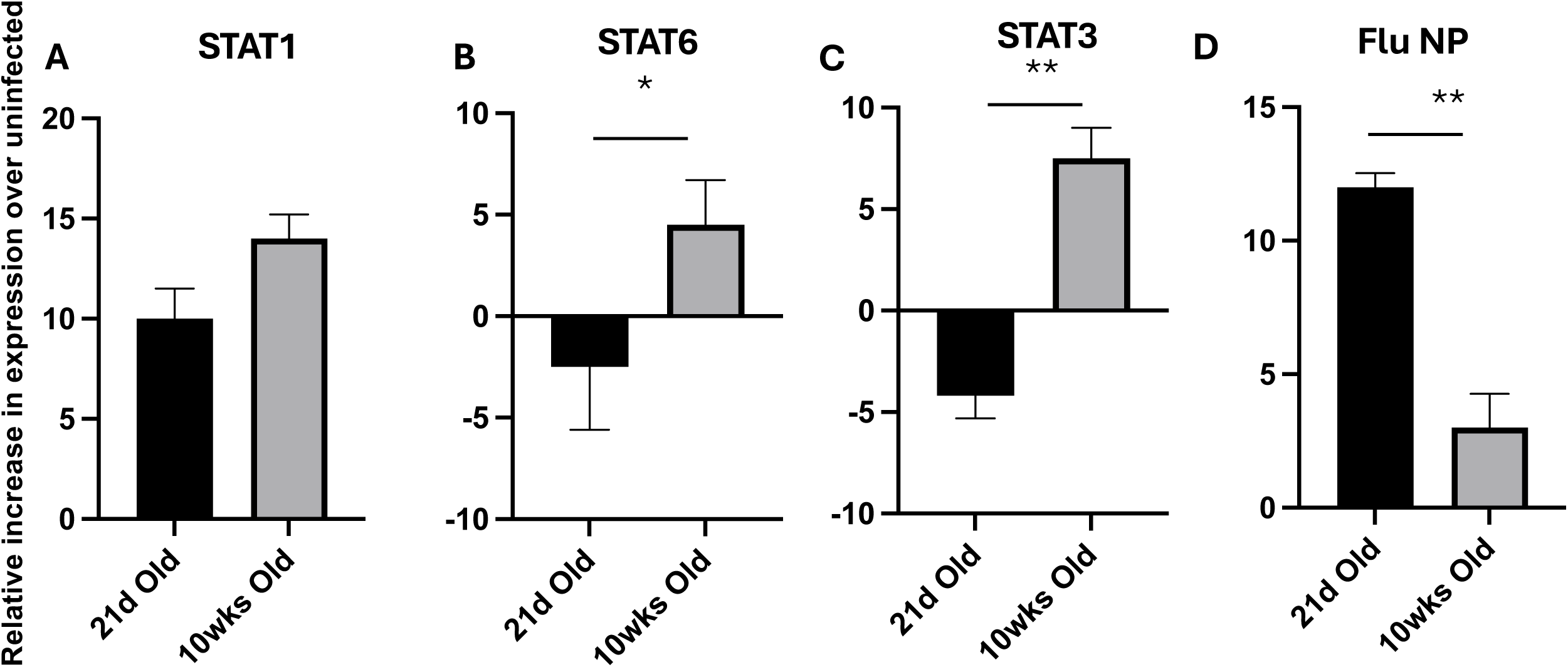
Divergent transcription factors associated with IFNγ production correlate with viral infection. We tested macrophages/monocytes harvested at day 5 post-infection for transcript expression of **(A)** STAT1, **(B)** STAT 6, and **(C)** STAT3 by qRT-PCR and expressed them as increases (or decreases) over age match uninfected controls. **(D)** Viral burdens in macrophages were also assessed in both populations by qRT-PCR and calculated as fold increases over uninfected controls (CT values of which were 39-40 for all tested). * p-value <0.05 or ** p-value < 0.001.

Since viral infection of monocytes and macrophages could impede their anti-viral functions, we next assessed the relative fold expression of influenza NP antigen by qRT-PCR at 5 DPI post-sublethal infection (Fig 5D). Of interest, we found that 21d old mice had a significantly higher viral transcript expression in their monocytes/macrophages then adult mice.

### Divergent balance of macrophage phenotypes in the lungs

To determine the phenotypes of macrophages/monocytes in young mice or adult comparators, we used flow cytometry with differential markers to determine the absolute numbers of each population throughout infection. 10wk old mice were characterized by an early influx of monocytes likely differentiating into inflammatory macrophages (Fig 6A-B). In fact, monocytes in the young seemed very delayed in infection and did not reach adult levels until day 7 post-infection which the innate immune system is often handing off to the adaptive. Interstitial macrophages and alveolar macrophages, both often resident cell types and associated with repair or inflammatory dampening were also minimally expansive in 21d old mice than older ones (Fig 6C-D).

**Figure 6.**
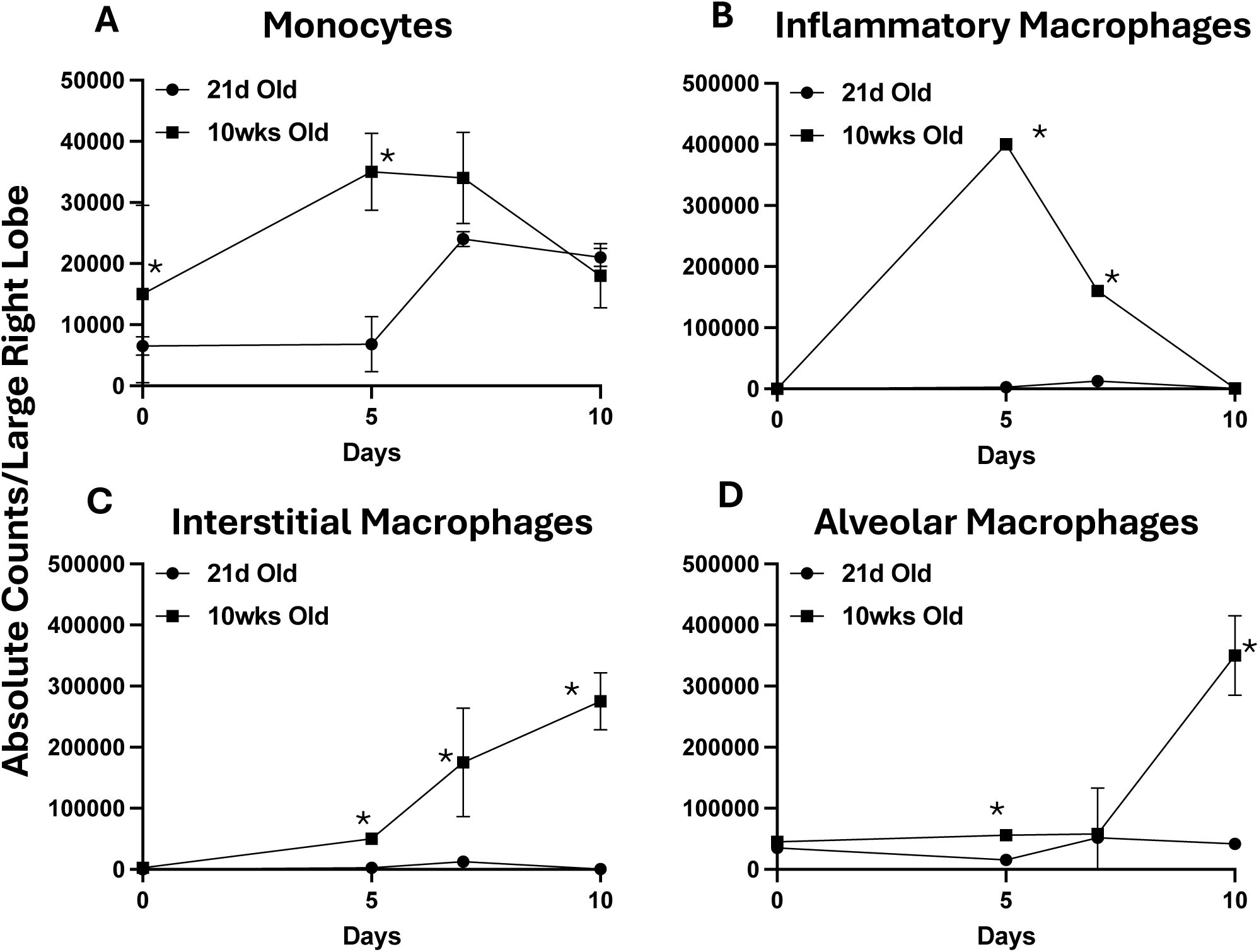
Differences macrophage or monocyte populations occur in the young and older mice during infection. We calculated the absolute counts of **(A)** monocytes, **(B)** inflammatory macrophage, **(C)** interstitial macrophages, and **(D)** alveolar macrophages from whole lungs at each time-point using flow cytometry. Markers for each population used were previous cited. * p-value <0.05

### Lung fibroblasts are master regulators of early innate responses in influenza but appear hampered in 21d old mice

We next isolated lung fibroblasts from uninfected mice and cultured them in tissue culture media. We then infected with H1N1 A/PR/8/34 these cells for 3 days and then determined the level of infection by intercellular staining. We found that lung fibroblasts form 21d old mice were strangely much easier to detect virus in than adults (Fig 7A). We next isolated fibroblasts at day 3 post-infection in 21d old and 10wk old mice and found that like fig 7A, fibroblasts in the younger mice appeared to be more condusive to infection with the virus (Fig 7B). We next determined the level of CCL2 expression since we saw such little monocyte recruitment in the young and found much less expression than older mice (Fig 7C).

**Figure 7.**
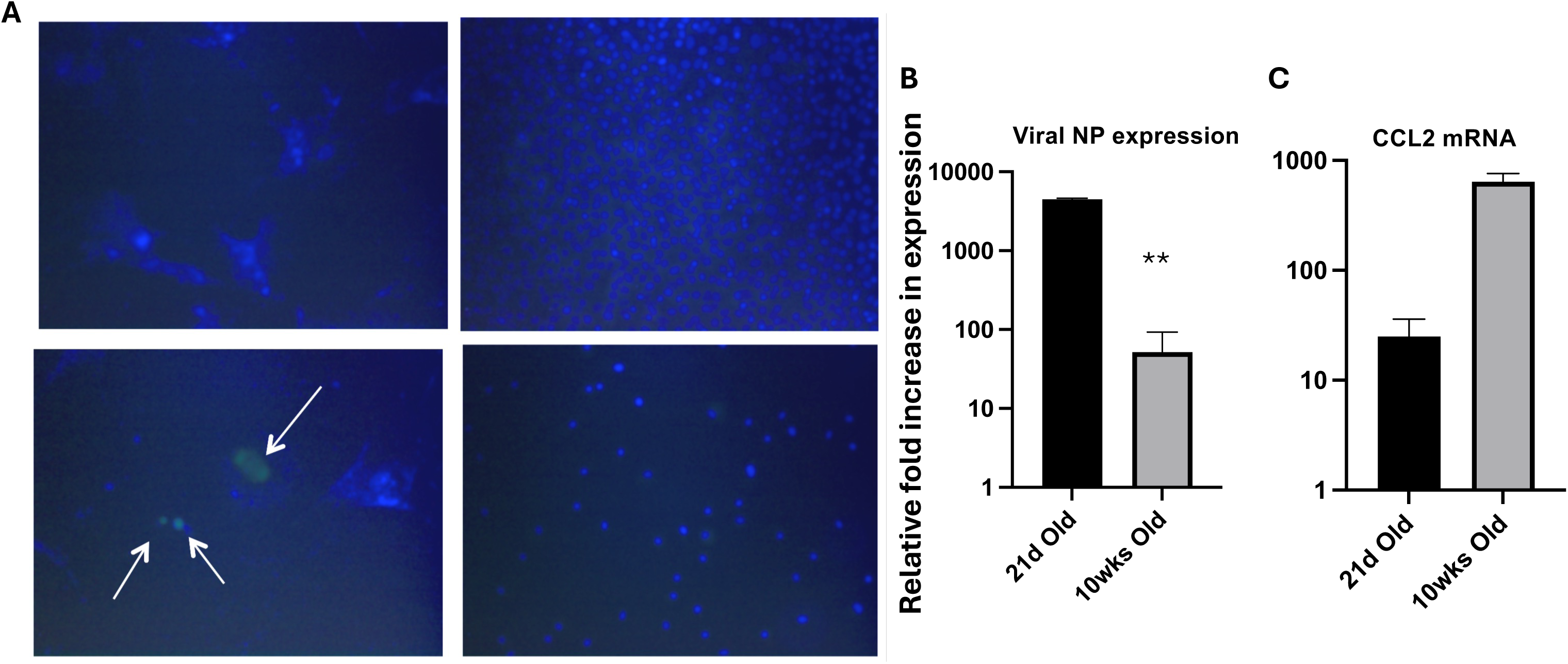
Disrupted fibroblast function correlates with impaired monocyte/macrophage recruitment. (A) We isolated fibroblasts from the lungs using tissue culture plate adherence and methods established in the methods section. We then infected fibroblasts with an MOI of 1 and fixed and intracellularly stained for influenza using a pan influenza A antibody. Green indicates cells infected with influenza while Blue indicated Dapi nuclear staining. A similar isolation technique was used with lungs from 5 day post-infected mice and levels of **(B)** influenza infection or **(C)** CCL2 transcript expression were determined by qRT-PCR immediately after plate attachment. In controls (not shown) staining the populations for macrophage markers or epithelial markers and given their morphology, we are reasonably assured that they were fibroblasts.

## Discussion

We sought here to examine whether key innate immunological limitations in young children due to immune maturation might be associated with the higher level of morbidity during respiratory viral infections. In our prior work, we found lower CD4 T-cell survival and function in young mice with this lack of CD4 T-cell signaling likely lowering feedback to macrophages to down regulate their responses at the innate to adaptive immune switch (27). While modeling young children in mice is often difficult we believe that our findings might relate to severe infection in very young children who do not control their virus well. However, it is well known that young children have lower innate immune response capacities than older peers (32–34). In our current study, we highlight key differences in the early innate immune response of young children especially as they relate to limited antiviral infections and key differences in macrophage phenotypes that respond to lung infection. Clearly, monocyte and macrophage limitations in antiviral responses continue from the more severe dysfunction common in neonates. Key differences in viral permissiveness, lower cytokine functions, and differences in macrophage populations occur between the young and adults.

In adult influenza infections, the level of pathology is related to the antiviral response as a “cytokine storm” perpetuates tissue histopathology irrespective of viral replication (35). Additionally, inflammatory monocytes are also known to increase the level of inflammatory cytokines in the lungs also contributing to the level of immunopathology. Here, we show that the morbidity in young mice may derive less from an over-response of the immune system and possibly from an under-response.

We previously found limitations in IFNγ and IL-10 secretion from CD4 T-cells affected both their survival and the level of immunopathology (27). We know from neonatal studies that STAT4 may be epigenetically regulated and thus an intrinsic inability to secrete IFNγ occurs in conjunction with an inability of antigen presenting cells to adequately stimulate CD4 T-cells (36). However, young mice appear to outgrow limitations in APC stimulatory functions and the limitations toward IFNγ secretion in CD4 T-cells appears downstream of STAT4 in the ERM transcription. While we detect similar lower IFNγ transcription in monocytes/macrophages clearly contribute to the lower tissue protein levels of this cytokine, what mechanism(s) causing lower IFNγ in this population of cells is/are currently unknown. Furthermore, the contributions of infection in these cells and their ability to secrete cytokines needs to be explored. Blocks could occur from a lack of early IFNγ signaling to macrophages from a lack of monocyte recruitment or whether other innate cells such as NK cells are also limited in young mice or if the limitation in secretion derives from other intrinsic blocks are all possible. Importantly, a lack of IFNγ may contribute to the unbalanced macrophage response observed in young mice that is dominated by the classical activation pathways. Whether a lack of IFNγ in young children is biologically beneficial because it limits immunopathology, especially as this population has reduced regulatory T-cells, is a matter of debate. However, lacking this cell clearly has important ramifications for limiting the antiviral response and controlling viral replication earlier in infection.

Appreciation for tissue resident cell populations and viral control is ever increasing in the immunology sphere. In our older mice, we found coordination between multiple macrophage phenotypes to both control infection and likely to control histopathology. Furthermore, CCR2+ monocytes are also important for recruitment of neutrophils and thus further recruitment of inflammatory monocytes. Here, we found the resident monocyte population non-responsive to early viral infection as the numbers between pre-infection and post-infection did not change. These data are different from another study that found inflammatory monocytes appeared to be associated with worse lung pathology and morbidity (37) however that study used older young mice (4wks) and a less pathogenic virus (H3N2 in mice). Cell apoptosis could also account for a lack of monocyte numbers in the lungs. Furthermore, there is a strong association between monocytes and CD4 T-cells with both populations able to influence the activity of the other. Thus, some of our lack of CD4 T-cell recruitment to the lungs from prior studies could be derived from low monocyte activity, or *vice versa*, a lack of very early CD4 T-cell infiltration in the lungs could influence monocyte recruitment.

A high level of infection of macrophages and permissive infection of fibroblasts is of interest especially as we do not detect differences in viral titers when evaluating tissue viral burdens for infectious virions (TCID_50_ assay) early in infection. These data could imply that much of the viral replication occurring in cells is abortive of fibroblasts. We also do not yet know the mechanisms for why fibroblasts appear more permissive to infection in younger mice and is something we will peruse in future studies.

In summary, young children suffer high levels of morbidity and mortality and do not usually have the compounding co-morbidities of the elderly who also frequently suffer during infection. The innate immune system is very important during early viral infections and thus limited monocyte/macrophage antiviral responses could be key to the high morbidity levels when these populations are infected with other viral or bacterial infections. The effect of immunological maturation is not widely appreciated in immune development of young children and thus vaccination or pathogen therapies would be best served to take these limitations into account. Children grow quickly especially in the very young and growth in the immune response is not always appreciated. A neonate’s immune response may widely differ from an infant, which may differ from toddlers who may also differ from older children and especially adults.

## Acknowledgements

DSV. and DDV. performed all assays and some data analysis. DDV. designed, performed assays, analyzed data, and wrote the manuscript.

## References

1. Disease NCfI. 2006. www.cdcgov.

2. CDC. 2006. National Center for Infectious Disease. www.cdcgov.

3. Lee HH, Hoeman CM, Hardaway JC, Guloglu FB, Ellis JS, Jain R, Divekar R, Tartar DM, Haymaker CL, Zaghouani H. 2008. Delayed maturation of an IL-12-producing dendritic cell subset explains the early Th2 bias in neonatal immunity. J Exp Med 205:2269–80.

4. Zaghouani H, Hoeman CM, Adkins B. 2009. Neonatal immunity: faulty T-helpers and the shortcomings of dendritic cells. Trends Immunol 30:585–91.

5. Welliver TP, Garofalo RP, Hosakote Y, Hintz KH, Avendano L, Sanchez K, Velozo L, Jafri H, Chavez-Bueno S, Ogra PL, McKinney L, Reed JL, Welliver RC, Sr. 2007. Severe human lower respiratory tract illness caused by respiratory syncytial virus and influenza virus is characterized by the absence of pulmonary cytotoxic lymphocyte responses. J Infect Dis 195:1126–36.

6. Welliver TP RJ, Welliver RC Sr. 2008. Respiratory syncytial virus and influenza virus infections: observations from tissues of fatal infant cases. Pediatric Infectious Diseases Journal 27:S92–S96.

7. Belnoue E, Fontannaz-Bozzotti P, Grillet S, Lambert PH, Siegrist CA. 2007. Protracted course of lymphocytic choriomeningitis virus WE infection in early life: induction but limited expansion of CD8+ effector T cells and absence of memory CD8+ T cells. J Virol 81:7338–50.

8. McCullers JA. 2006. Insights into the interaction between influenza virus and pneumococcus. Clin Microbiol Rev 19:571–82.

9. Short KR, Habets MN, Hermans PW, Diavatopoulos DA. 2012. Interactions between Streptococcus pneumoniae and influenza virus: a mutually beneficial relationship? Future Microbiol 7:609–24.

10. Nakamura S, Davis KM, Weiser JN. 2011. Synergistic stimulation of type I interferons during influenza virus coinfection promotes Streptococcus pneumoniae colonization in mice. J Clin Invest 121:3657–65.

11. Gordon SB, Read RC. 2002. Macrophage defences against respiratory tract infections. Br Med Bull 61:45–61.

12. Cardani A, Boulton A, Kim TS, Braciale TJ. 2017. Alveolar Macrophages Prevent Lethal Influenza Pneumonia By Inhibiting Infection Of Type-1 Alveolar Epithelial Cells. PLoS Pathog 13:e1006140.

13. Kaur M, Bell T, Salek-Ardakani S, Hussell T. 2015. Macrophage adaptation in airway inflammatory resolution. Eur Respir Rev 24:510–5.

14. Schneider C, Nobs SP, Heer AK, Kurrer M, Klinke G, van Rooijen N, Vogel J, Kopf M. 2014. Alveolar macrophages are essential for protection from respiratory failure and associated morbidity following influenza virus infection. PLoS Pathog 10:e1004053.

15. Lin SJ, Lo M, Kuo RL, Shih SR, Ojcius DM, Lu J, Lee CK, Chen HC, Lin MY, Leu CM, Lin CN, Tsai CH. 2014. The pathological effects of CCR2+ inflammatory monocytes are amplified by an IFNAR1-triggered chemokine feedback loop in highly pathogenic influenza infection. J Biomed Sci 21:99.

16. Lin KL, Suzuki Y, Nakano H, Ramsburg E, Gunn MD. 2008. CCR2+ monocyte-derived dendritic cells and exudate macrophages produce influenza-induced pulmonary immune pathology and mortality. J Immunol 180:2562–72.

17. Friesenhagen J, Boergeling Y, Hrincius E, Ludwig S, Roth J, Viemann D. 2012. Highly pathogenic avian influenza viruses inhibit effective immune responses of human blood-derived macrophages. J Leukoc Biol 92:11–20.

18. Laskin DL, Weinberger B, Laskin JD. 2001. Functional heterogeneity in liver and lung macrophages. J Leukoc Biol 70:163–70.

19. Blumenthal RL, Campbell DE, Hwang P, DeKruyff RH, Frankel LR, Umetsu DT. 2001. Human alveolar macrophages induce functional inactivation in antigen-specific CD4 T cells. J Allergy Clin Immunol 107:258–64.

20. Thepen T, Van Rooijen N, Kraal G. 1989. Alveolar macrophage elimination in vivo is associated with an increase in pulmonary immune response in mice. J Exp Med 170:499–509.

21. Hamilton-Easton A, Eichelberger M. 1995. Virus-specific antigen presentation by different subsets of cells from lung and mediastinal lymph node tissues of influenza virus-infected mice. J Virol 69:6359–66.

22. Yu WC, Chan RW, Wang J, Travanty EA, Nicholls JM, Peiris JS, Mason RJ, Chan MC. 2011. Viral replication and innate host responses in primary human alveolar epithelial cells and alveolar macrophages infected with influenza H5N1 and H1N1 viruses. J Virol 85:6844–55.

23. Short KR, Brooks AG, Reading PC, Londrigan SL. 2012. The fate of influenza A virus after infection of human macrophages and dendritic cells. J Gen Virol 93:2315–25.

24. Gordon S. 2003. Alternative activation of macrophages. Nat Rev Immunol 3:23–35.

25. Leopold Wager CM, Wormley FL, Jr. 2014. Classical versus alternative macrophage activation: the Ying and the Yang in host defense against pulmonary fungal infections. Mucosal Immunol 7:1023–35.

26. Verhoeven D, Perry S. 2016. Differential mucosal IL-10-induced immunoregulation of innate immune responses occurs in influenza infected infants/toddlers and adults. Immunol Cell Biol doi:10.1038/icb.2016.91.

27. Verhoeven D, Perry S, Pryharski K. 2016. Control of influenza infection is impaired by diminished interferon-gamma secretion by CD4 T cell in the lungs of toddlers. J Leukoc Biol doi:10.1189/jlb.4A1014-497RR.

28. Verhoeven D, Teijaro JR, Farber DL. 2009. Pulse-oximetry accurately predicts lung pathology and the immune response during influenza infection. Virology 390:151–6.

29. Teijaro JR, Verhoeven D, Page CA, Turner D, Farber DL. Memory CD4 T cells direct protective responses to influenza virus in the lungs through helper-independent mechanisms. J Virol 84:9217–26.

30. Chandran SS, Verhoeven D, Teijaro JR, Fenton MJ, Farber DL. 2009. TLR2 engagement on dendritic cells promotes high frequency effector and memory CD4 T cell responses. J Immunol 183:7832–41.

31. Luo H, Wang D, Che HL, Zhao Y, Jin H. 2012. Pathological observations of lung inflammation after administration of IP-10 in influenza virus- and respiratory syncytial virus-infected mice. Exp Ther Med 3:76–79.

32. Verhoeven D. 2019. Immunometabolism and innate immunity in the context of immunological maturation and respiratory pathogens in young children. J Leukoc Biol 106:301–308.

33. Verhoeven D. 2019. Influence of Immunological Maturity on Respiratory Syncytial Virus-Induced Morbidity in Young Children. Viral Immunol 32:76–83.

34. Cole SL, Dunning J, Kok WL, Benam KH, Benlahrech A, Repapi E, Martinez FO, Drumright L, Powell TJ, Bennett M, Elderfield R, Thomas C, investigators M, Dong T, McCauley J, Liew FY, Taylor S, Zambon M, Barclay W, Cerundolo V, Openshaw PJ, McMichael AJ, Ho LP. 2017. M1- like monocytes are a major immunological determinant of severity in previously healthy adults with life-threatening influenza. JCI Insight 2:e91868.

35. Gu Y, Zuo X, Zhang S, Ouyang Z, Jiang S, Wang F, Wang G. 2021. The Mechanism behind Influenza Virus Cytokine Storm. Viruses 13.

36. Tang ML, Kemp AS, Thorburn J, Hill DJ. 1994. Reduced interferon-gamma secretion in neonates and subsequent atopy. Lancet 344:983–5.

37. Coates BM, Staricha KL, Koch CM, Cheng Y, Shumaker DK, Budinger GRS, Perlman H, Misharin AV, Ridge KM. 2018. Inflammatory Monocytes Drive Influenza A Virus-Mediated Lung Injury in Juvenile Mice. J Immunol 200:2391–2404.

